# Exosome Release Promotes Inflammatory Resolution in Activated and Aged Microglia

**DOI:** 10.1101/423558

**Authors:** Joe C. Udeochu, Cesar Sanchez-Diaz, Alvan Cai, Anna Jovicic, Saul A. Villeda

**Affiliations:** Department of Anatomy, University of California San Francisco, San Francisco, California 94143, USA; The Eli and Edythe Broad Center for Regeneration Medicine and Stem Cell Research, San Francisco, California 94143, USA; Biomedical Sciences Graduate Program, University of California San Francisco, San Francisco, California 94143, USA; Department of Genetics, Stanford University, Stanford, California 94305, USA; Department of Physical Therapy and Rehabilitation Science, University of California San Francisco, San Francisco, California 94143, USA

## Abstract

Proper regulation of inflammatory responses is critical for effective function of microglia, in both physiological and disease states. While the mechanisms that drive microglia activation are well characterized, the pathways leading to inflammatory resolution and immune homeostasis have yet to be fully elucidated. Using RNA interference, pharmacological inhibition and genetic knockout mouse model approaches, we show that exosome release promotes immune homeostasis in activated and aged microglia. We demonstrate that induction of anti-inflammatory pathways enhances release of exosomes containing immune proteins and microRNAs. Functionally, inhibition of exosome release alters trafficking of exosome cargo, such as miR-155, in activated microglia resulting in increased cellular retention of these cargo molecules. Concordantly, we identify increased miR-155 activity leading to sustained activation of pro-inflammatory pathways as a potential mechanism underlying impaired inflammatory resolution due to inhibition of microglia exosome release. Similarly, inhibition of augmented exosome release in aged microglia exacerbates inflammatory activation, demonstrating conservation of the immune modulatory effects of exosome release in microglia. Taken together, our study identifies exosomes as novel components of an anti-inflammatory mechanism utilized by activated and aged microglia to restore immune homeostasis.

## Introduction

As the tissue macrophages of the brain, microglia activity is strongly tailored to meet the specialized needs of their environment (Kettenmann et al., 2011; Nimmerjahn et al., 2005). Increased brain inflammatory milieu during aging and disease, however, drives transformation of microglia from a homeostatic to an activated state (Hickman et al., 2013; Prinz et al., 2011; Udeochu et al., 2016). Numerous factors including increased cytokine production, cellular senescence, and loss of neuronal ‘off’ signals have been implicated in driving microglia inflammatory transformation (Wong, 2013; Ye and Johnson, 1999). However, much still remains to be learned about mechanisms that counterbalance the effects of inflammation to facilitate immune homeostasis in activated aged and diseased microglia.

The endolysosomal network is central for maintenance of cell homeostasis, through trafficking, signaling, and degradation of both endocytic and biosynthetic cargoes (Klumperman and Raposo, 2014). Exosomes, a subset of endosomal-derived extracellular vesicles, have emerged as critical regulators of various aspects of peripheral immune cell function including inflammatory activation and suppression (Lindenbergh and Stoorvogel, 2018; Robbins and Morelli, 2014). Moreover, mutations in regulator of exosome release, Rab27a, cause Griscelli syndrome, which is characterized by peripheral macrophage hyperactivation and cellular infiltration of the brain (Menasche et al., 2000; Sarper et al., 2002). Recent studies have demonstrated microglia uptake of exosomes produced by neurons and other glia (Fitzner et al., 2011; Fruhbeis et al., 2013), and regulation of neuropeptide catabolism by microglia exosomes (Potolicchio et al., 2005). Exosome involvement in microglia inflammation is, however, unclear. To address this, we utilized molecular and genetic techniques to assess exosomal regulation of inflammation in cultured, adult brain and aged microglia. Specifically, we studied the effects of microglia activation on exosome biogenesis, manipulated exosome biogenesis experimentally, and determined a molecular mechanism mediating exosomal regulation of immune homeostasis in activated microglia.

## Results

### Microglia produce canonical exosomes that contain cell type-specific factors

To begin our study, we characterized exosome production in primary microglia and BV2 immortalized microglia cell line cultured under serum free conditions. Secreted exosomes were isolated from media by standard? filtration and differential ultracentrifugation (Thery et al., 2006). Analysis of physical characteristics and protein content revealed enrichment of exosomes in our preparations. Specifically, scanning electron microscopy and nanoparticle tracking analysis (NTA) showed that microglia exosomes exhibit stereotypical donut-shaped morphology and were on average 30-120nm in size (Figures 1A and B). Detection of tetraspanin proteins, CD9 and CD63, and endolysosomal protein, LAMP2, further confirmed exosome enrichment (Figure 1C). We next tested the involvement of ceramide synthase pathway and Rab GTPases in microglia exosome biogenesis by inhibiting neutral sphingomyelinase 2 (nSMase2) and Rab27a, respectively (Ostrowski et al., 2010; Trajkovic et al., 2008). Transduction of BV2 microglia with lentiviral particles expressing Rab27a targeting (KD) short hairpins RNAs (shRNAs) led to efficient Rab27a knockdown (Figure 1D) and reduced exosome secretion into media compared to cells expressing scrambled (scr) control sequences (Figures 1E and 1F). Similarly, treatment with the neutral sphingomyelinase inhibitor, GW4869, reduced microglia exosome production (Figure 1G).

**Figure 1.**
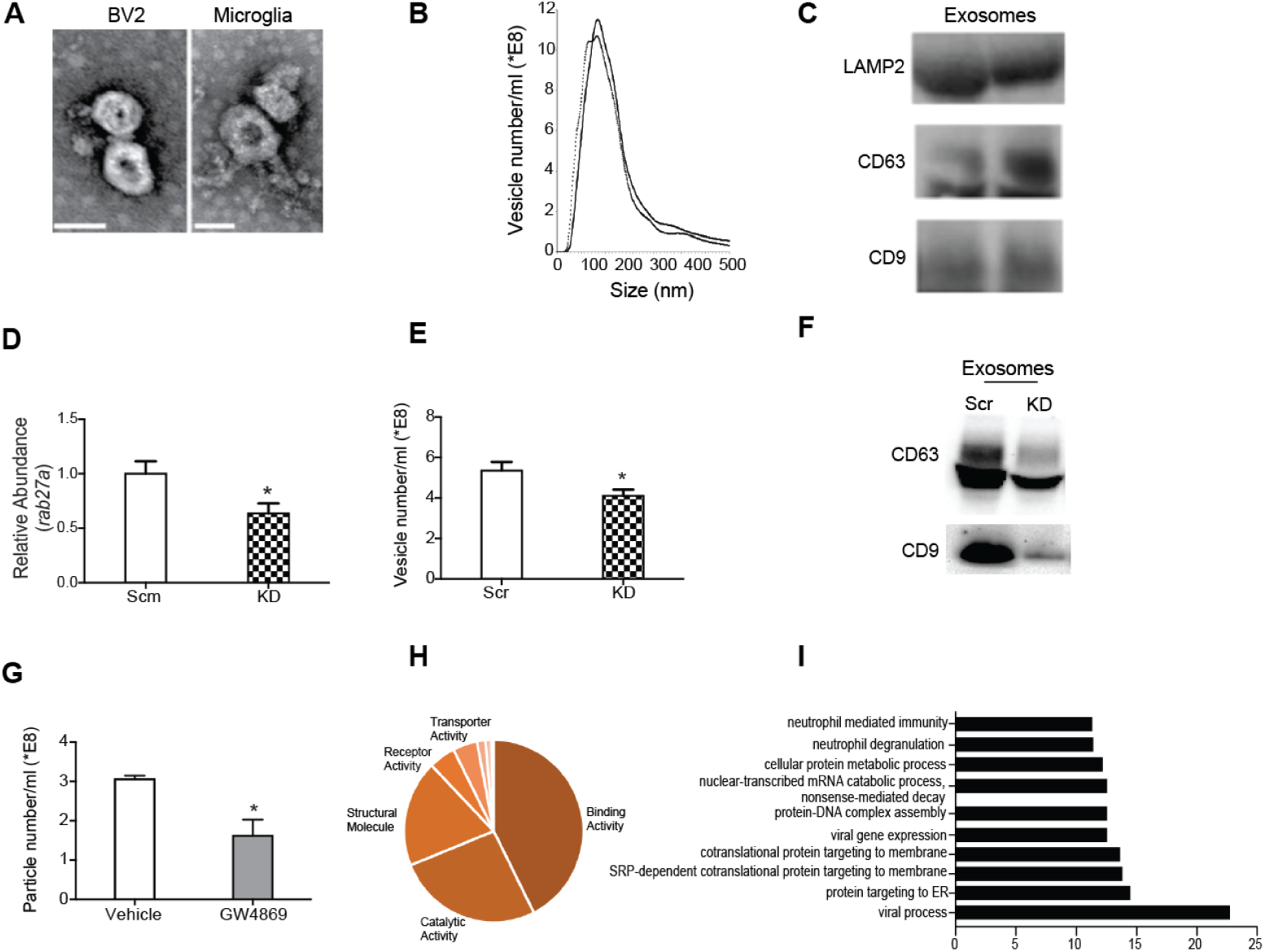
Microglia inflammatory activation regulates expression of exosome regulatory genes. (A) Representative electron micrographs showing prototypical cup-shaped morphology and size of BV2 and primary microglia exosomes (Scale bar = 50nm). (B) Nanoparticle tracking analysis of replicate exosome preps shows heterogeneous vesicle size, with most abundant vesicles ranging in size from 50 – 130nm. (C) Western blot analysis of exosome lysates showing expression of tetraspanin (CD9 and CD63) and endocytic (LAMP2) proteins. (D) Lentiviral mediated transduction of rab27a shRNA (kd) leads to reducted rab27a transcript levels compared to scramble (scm) shRNA control BV2 microglia. (E,F) Rab27a knockdown results in decreased exosome production as determined by as determined by nanoparticle tracking (E) and Western blot (F) analyses. (G) Inhibition of neutral sphingomyelinase 2 (nSMase2) by GW4869 results in reduced microglia exosome production. (H) Pie chart representation of heterogeneity in BV2 microglia exosome protein content classified by molecular function. (I) Bar graph showing over-represented pathways detected in microglia exosome proteins. All data represented as mean ± SEM; ^∗^P<0.05, t-test (D, E, G), Bonferroni correction for multiple testing used to identify overrepresented proteins, FDR <0.05 (I). Scale bar = 50 nm

To elucidate the signaling potential of microglial exosomes, we characterized the protein and miRNA contents of exosomes produced by BV2 microglia. Mass spectrometry and miRNA array analyses revealed that microglia exosomes contain heterogeneous cargo, as has been reported for exosomes from other cell types (Kowal et al., 2016; Squadrito et al., 2014) (Figure 1H, Supplemental Table 1 and 2). We identified canonical exosome content such ALIX and MFGE8 proteins, as well as cargo unique to microglia and myeloid cells like mannose receptor, macrophage inhibitory protein, miR-146a, and miR-150 (Supplemental Table 1 and 2). Gene ontology analysis revealed that microglia exosomes were enriched in proteins involved in biological processes such as viral gene expression, protein trafficking, nucleic acid catabolism and degranulation (Figure 1I). The diversity of biological pathways represented in microglia exosome cargo suggested a potential role in regulating cellular responses and function.

### Microglia activation leads to recruitment of pro-inflammatory molecules into exosomes

To explore exosome involvement in microglia inflammatory response, we next assessed the effects of activation on the expression of exosome regulatory genes, rab27a and nSMase2. Quantitative real time PCR analyses of microglia stimulated with pro-inflammatory agents - interferon gamma (IFNγ), lipopolysaccharide (LPS), or polyribocytidilic acid (poly I:C), revealed activation induced changes in Rab27a and nSMase2 expression (Figure 2A and S1). Overall, pro-inflammatory activation suppressed microglia nSMase2 expression. On the other hand, Rab27a expression was more dynamically regulated in activated microglia. Strikingly, we note that kinetics of Rab27a expression mirrored interleukin 4 (IL-4) under all three activation paradigms. Consequently, we tested the possibility that IL-4 might coordinate Rab27a expression and show that recombinant IL-4 stimulation significantly increases mi croglia Rab27a (Figure 2B). These findings show that exosome regulatory genes are responsive to microglia activation, and suggest an anti-inflammatory function for exosome release in acutely activated microglia.

**Figure 2.**
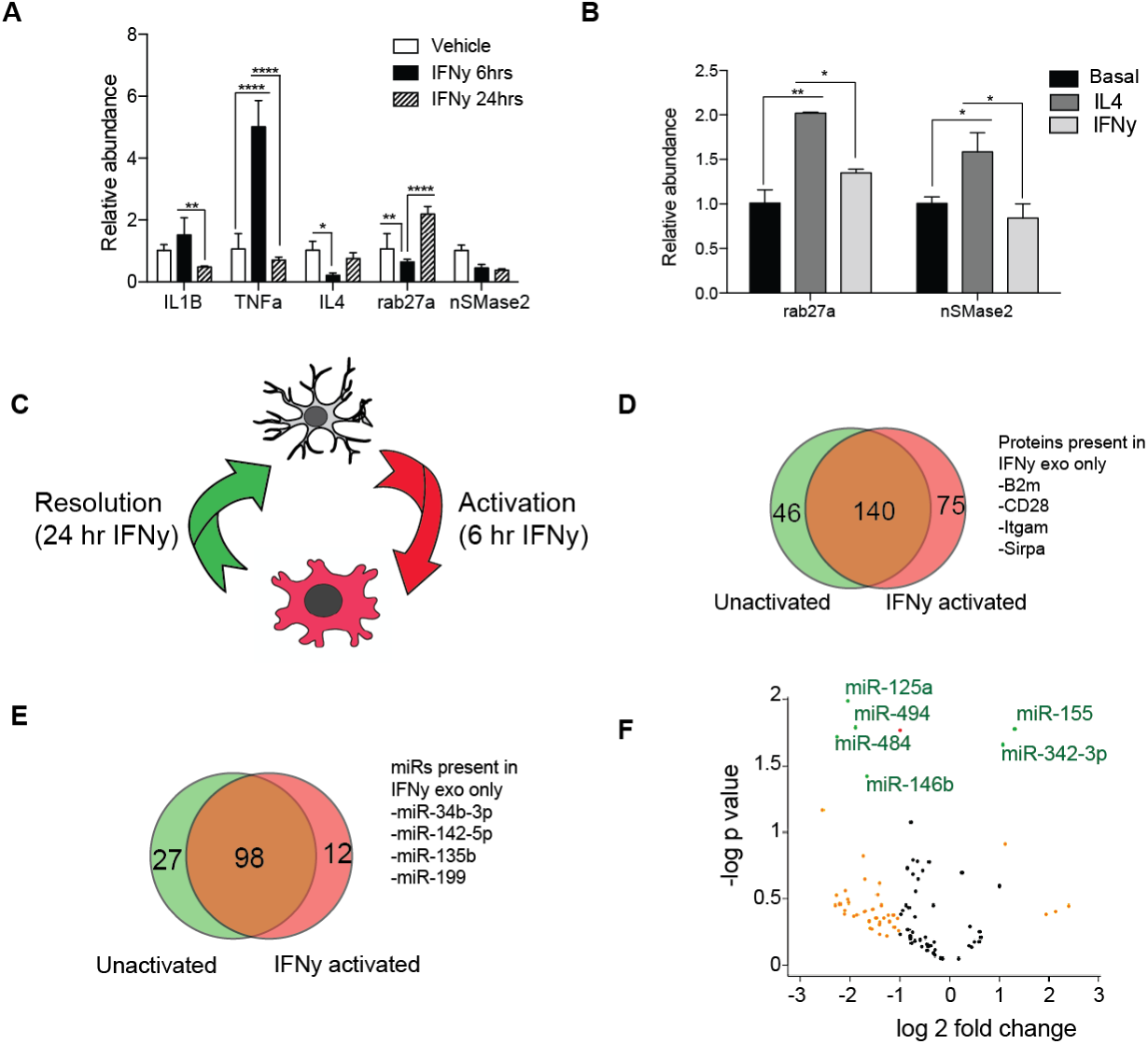
Inflammation regulates exosome production and cargo in microglia. (A) Quantitative real-time PCR results of interferon gamma (IFNψ) stimulated primary microglia showing expression of inflammatory cytokines (IL1β, TNFα, and IL4) and exosome regulatory genes (nSMase2 and rab27a). (B) Acute interleukin-4 stimulation significantly upregulates rab27a in BV2 microglia. (C) Schematic demonstrating model for IFNγ-induced inflammatory activation and resolution in microglia. (D-E) Comparison of protein (D) and microRNA (E) contents of exosomes derived from activated and unactivated microglia during resolution of IFNγ inflammation. (F) Volcano plot showing IFNγ induced enrichment of immune related miRNAs, miR-155 and miR-342-3p, in IFN exosomes compared unactivated exomes (yellow = 2 fold change, red = p < 0.05, green = 2 fold change and p < 0.05). All data represented as mean ± SEM; ^∗^P<0.05; ^∗∗^P<0.01; ^∗∗∗∗^P<0.0001, one-way ANOVA (A), t-test (B, F).

Further inspection of cytokine gene expression kinetics revealed that the 6 and 24 hour IFNγ stimulation timepoints represent two distinct phases of inflammation – activation and resolution phases, respectively (Figure 2C). Given that Rab27a is strongly upregulated during resolution and coincides with the repression of pro-inflammatory IL1b and TNFa, we next compared the protein and miRNA content of exosomes released by resolving and unstimulated BV2 microglia for differences in immune signatures. While exosomes from resolving and unstimulated microglia shared many proteins and miRNAs, we observed some activation induced differences in cargo (Figure 2F-2H). There was selective or increased recruitment of cargo implicated in immune cell activation, such as CD28, beta-2-microglobulin, miR-142-5p, and miR-155, (Ponomarev et al., 2013) into exosomes released by resolving microglia. This suggests that exosome cargo sorting in microglia is dynamic and partly influenced by microglia activation state.

### Exosome release is necessary for efficient resolution of inflammation in microglia

Given the observed enrichment of immune related miRNAs in IFNγ exosomes, we next assessed the consequences of inhibiting exosome biogenesis on the trafficking of these miRNAs. While abrogation of Rab27a had no effect on basal levels of miR-103 and miR-155, we observed increased cellular accumulation of miR-155 in resolving Rab27a KD BV2 compared to scr controls (Figure 3A). Similarly, GW4869 treatment increased cellular accumulation of miR-142-5p and miR-155 after IFNγ stimulation (Figure S2A). miR-155 activity amplifies interferon induced JAK/ STAT signaling in part by repressing inhibitors of immune response such as the suppressor of cytokine signaling (SOCS1) (Wang et al., 2010). Quantitative PCR analyses revealed significantly lower SOCS1 expression in Rab27a KD and GW4869 treated BV2 compared to respective controls, indicating increased miR-155 activity (Figures 3B and S2B). To determine if these observed changes in exosomal miR-155 trafficking resulted in broad inflammatory dysregulation in activated microglia, we measured the expression of various pro-inflammatory mediators of IFNγ signaling. Indeed, resolving Rab27a KD microglia expressed significantly higher levels of IL1β, myd88, and Stat1 compared to scr controls (Figure 3C). GW4869 treatment resulted in similar but milder effects on microglia activation, in line with data in Figure 2 showing more prominent induction of Rab27a compared to nSmase2 during resolution of IFNγ inflammation (Figure S2C). Concordantly, resolving Rab27a KD and GW4869-treated microglia exhibited increased Stat1 phosphorylation levels compared to controls, indicating sustained activation of JAK-STAT signaling (Figure 3D and S2D). These findings highlight a novel role of exosomes in microglia immune homeostasis, wherein increasing exosome release promotes clearance of immune miRNAs like miR-155 to allow inflammatory resolution.

**Figure 3.**
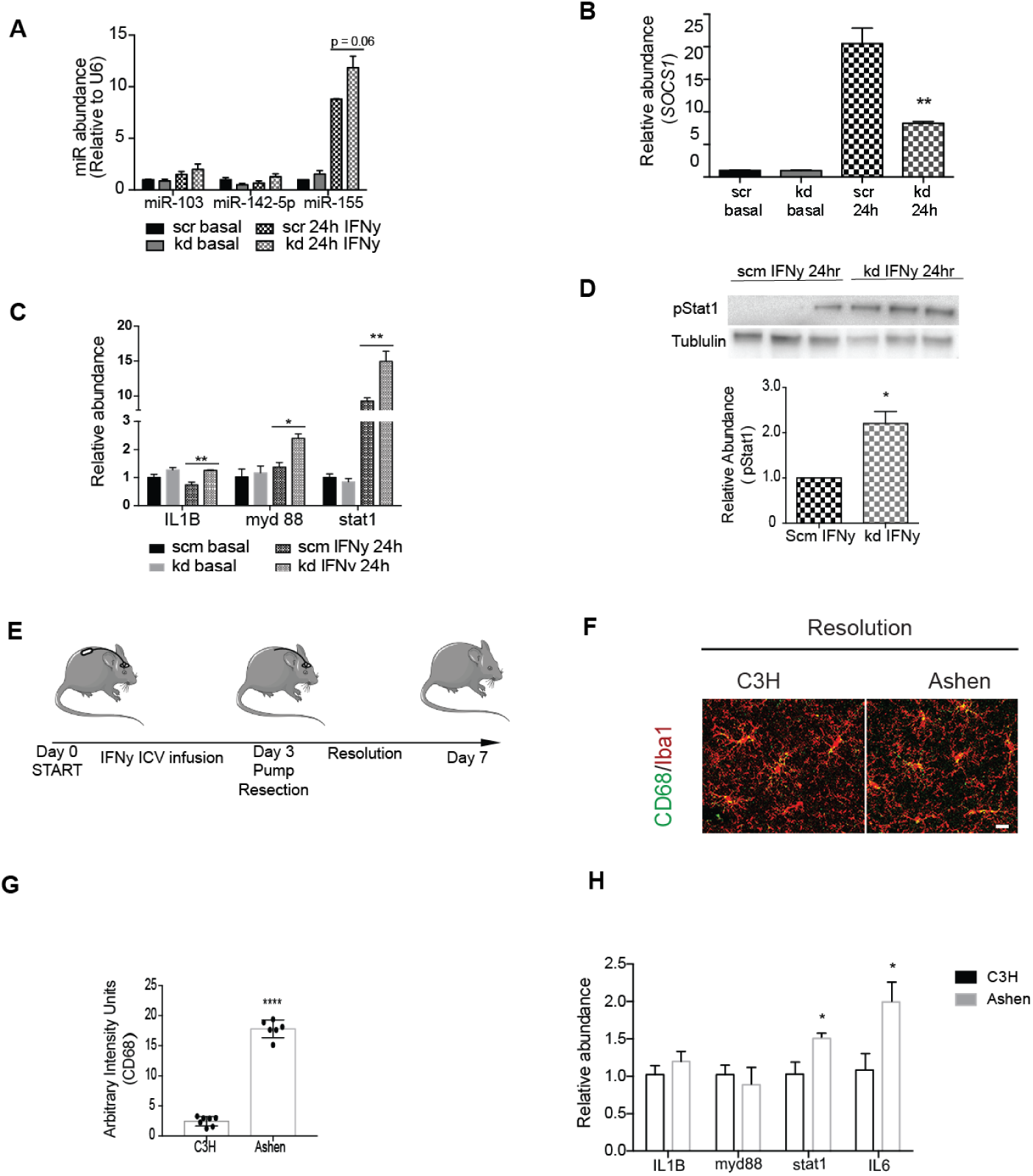
Inhibition of exosome productions alters miR-155 dynamics and impairs resolution of acute inflammation in cultured and adult brain microglia. (A) Quantitative PCR analyses of IFNγ exosome enriched miRNAs, showing increased cellular miR-155 levels in activated rab27a KD compared to scm BV2. (B) Corresponding decrease in expression of miR-155 target, suppressor of cytokinen signaling (SOCS1), in activated rab27a KD BV2. (C) Increased transcript levels of pro-inflammatory mediators IL1β, myd88 and Stat1 in rab27a KD compared to scm BV2 microglia. (D) Sustained STAT1 phosphorylation in rab27a KD compared scm BV2 microglia 24 hours post IFNγ activation. (E) Intracerebroventricular (ICV) osmotic infusion paradigm utilized to assess IFNγ induced inflammatory resolution in 3 month old wild type (C3H) and rab27a mutant (Ashen) adult brain microglia. (F) Representative confocal images showing higher CD68 immunoreactivity in Iba1 + microglia in Ashen compared to C3H brains, quantified in (G). (H) Quantitative PCR analyses showing increased Stat1 and IL6 transcript levels in Ashen compared to C3H brain homogenates, indicating impaired resolution of IFNγ inflammation. Data represented as mean ± SEM; ^∗^P<0.05; ^∗∗^P<0.01; unpaired t-test (A-D, G-H). Scale bar = 20um.

Next, we examined whether inhibiting exosome release in the adult brain would also impairs resolution of microglia inflammation after acute IFNγ stimulation. We compared microglia activation in wild type C3H mice and Ashen mice, which harbor a spontaneously generated point mutation in Rab27a that results in a hypomorphic allele. Under baseline conditions, no differences in microglia activation were observed between young (3 months) Ashen mice and C3H controls, as assessed by immunofluorescence analysis of the microglia-specific calcium binding adaptor protein, Iba1, and the activation marker, CD68, respectively (Figures S3A and B). To model resolution of IFNγ-mediated inflammation, we perfomed a modified intracerebroventricular (ICV) infusion paradigm on 3-month old C3H and Ashen brains. Mice received ICV IFNγ infusion, after which osmotic pumps were surgically resected and mice allowed four days of recovery (Figure 3E). Immunofluorescent analysis post recovery revealed low levels of CD68 immunoreactivity in C3H wild type microglia indicating effective inflammatory resolution of IFNγ-induced inflammation. Ashen microglia, however, maintained significantly higher CD68 expression indicating impaired resolution (Figure 3F and 3G). In addition, higher Stat1 and IL6 levels show sustained IFNγ activation in the resolving Ashen compared to C3H brains (Figure 3H). We also analyzed microglia activation immediately after three days IFNγ infusion (Figure S3C) and observed no differences in CD68 expression between Ashen and C3H microglia (Figures. S3D and E). Thus, the differences reported here between Ashen and C3H microglia during inflammatory resolution are unlikely due to differential IFNγ activation (Figures. S3D and E). Our findings demonstrate exosome release is required for efficient inflammatory resolution in acutely activated young adult brain microglia.

### Augmented exosome release modulates inflammatory activation in aged microglia

Microglia aging is accompanied by increased production of inflammatory molecules resulting in a chronic state of activation that drives morphological and functional changes in microglia (Mosher and Wyss-Coray, 2014; Sierra et al., 2007). Upregulation of interferon signaling and other antiviral response pathways has recently been identified as one of the transcriptional signatures associated with activation in aged microglia (Baruch et al., 2014; Grabert et al., 2016). Considering our findings implicating exosomes in interferon responses in acutely activated microglia, we examined the potential involvement of exosomes in aging-associated chronic microglia activation. We first compared the abundance of interstitial exosomes between young and aged brains, using a previously published protocol (Perez-Gonzalez et al., 2012). Interstitial exosomes were higher in aged compared to young brains, based on increased CD9 and CD63 immunolabeling and nanoparticle tracking analyses (Figures 4A-4C). To directly test the effects of aging on microglia exosome production, we utilized qPCR analyses of exosome regulatory genes in acutely isolated young and aged microglia. Higher transcript levels of IL1β and IL10 in aged microglia confirmed the activated state of aged microglia compared to young (Figure 4D). Interestingly, we also observed increased rab27a expression in aged microglia, indicating that microglia activation in the aged brain is accompanied by increased exosome release (Figure 4D). Furthermore, immunofluorescence analysis revealed higher CD63 levels in aged compared to young microglia (Figures 4E and 4F). These data demonstrate age-dependent alterations in microglia exosome production and suggest that the increase of exosome abundance in the aged brain derives, at least in part, from increased exosome release by aged microglia.

**Figure 4.**
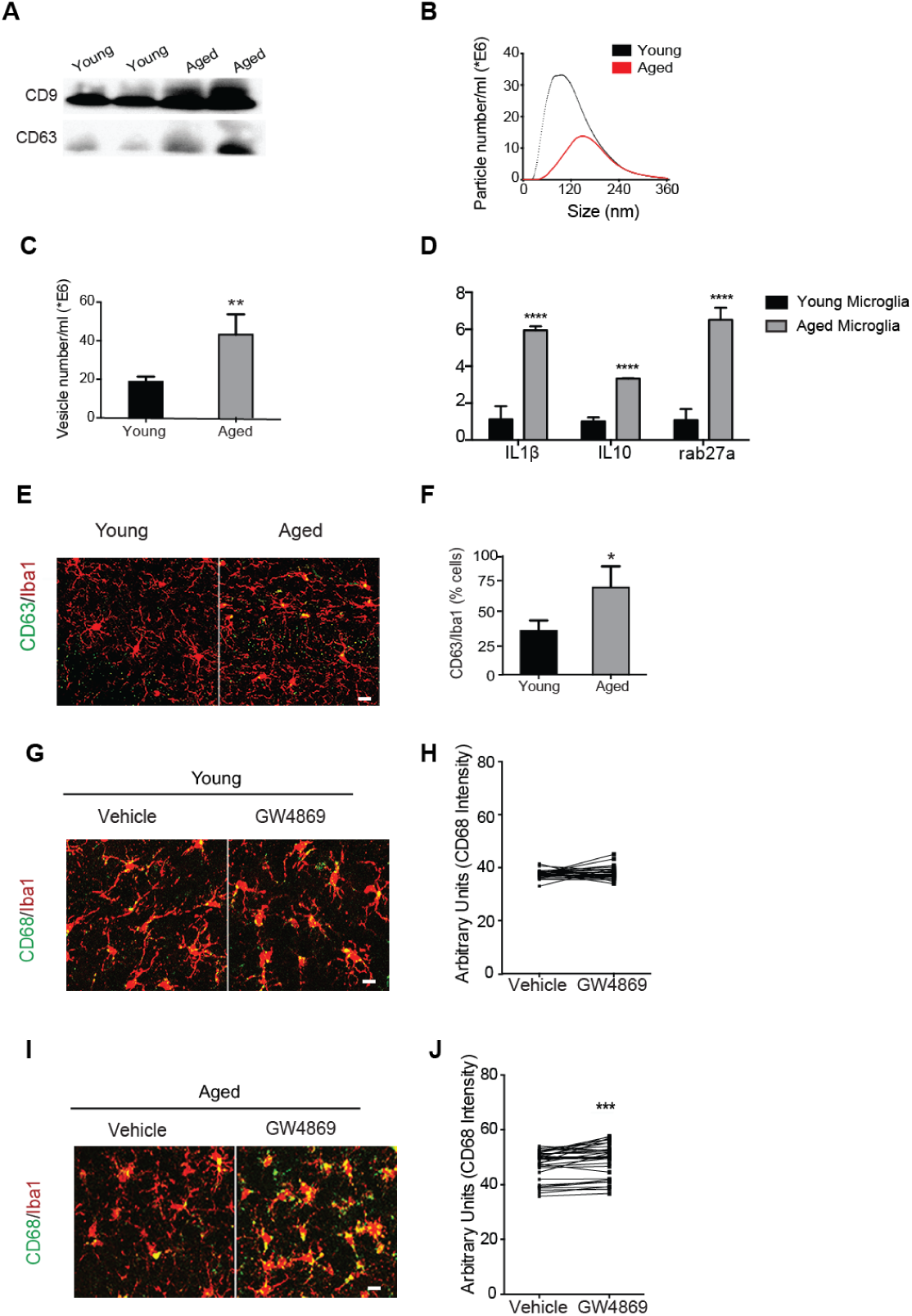
Increased exosome production modulates inflammatory activation in aged microglia. (A) Representative western blot probes for tetraspanin proteins, CD9 and CD63, in exosomes isolated from young (3 months) and aged (18 months) hemibrains. (B) Representative nanoparticle tracking traces of young and aged brain exosomes. (C) Nanoparticle tracking analyses showing significantly higher abundance of interstitial exosomes in aged compared to young brains. (D) Quantitative PCR of acutely isolated young and aged exosomes, showing upregulated expression of inflammatory (Il1B, Il10) and exosome regulatory (rab27a) genes. (E) Representative immunofluorescent images of microglia (Iba1) colabeled with exosome marker (CD63). (F) Quantification of hippocampal microglia showing aging-associated increase in microglial CD63. (G,I) Representative immunofluorescent images of vehicle and GW4869 injected hemibrains stained for microglia marker, Iba1, and activation marker, Cd68. (H) No difference in CD68 intensity in young vehicle and GW4869 hemibrains. (J) GW4869 treatment results in significantly higher CD68 expression compared to vehicle treated hemibrains in aged mice. Data represented as mean ± SEM; ^∗^P<0.05; ^∗∗^P<0.01; ^∗∗∗∗^P<0.0001; t-test (C, D, F, H, J). Scale bar = 20μm.

Lastly, we asked if augmented exosome production is involved in the regulation of inflammation in aged microglia. We performed stereotaxic injections of exosome inhibitor, GW4869, or vehicle into contralateral hippocampi of young (3 months) and aged (18 months) mice. Microglia activation was assessed one week post-injection by CD68 immunofluorescence analyses. No differences in CD68 expression were detected between GW4869 and vehicle injected conditions in the young brain (Figures 4G and 4H). In the aged brain, however, microglia CD68 expression was significantly higher in GW4869-compared to vehicle-injected hippocampi (Figures 4I and 4J). These data show that augmented exosome release is an important anti-inflammatory mechanism utilized by microglia to modulate aging-associated chronic inflammatory activation.

## Discussion

Microglia transform to an activated state in response to inflammatory challenge and changes in brain homeostasis during aging and disease. There is increasing evidence that sustained microglia activation can contribute to brain pathology and dysfunction under these conditions. A better understanding of microglia inflammatory processes can inform ongoing efforts to preserve and restore function in aged and diseased brains. Our study focuses not on drivers of microglia activation, but on mechanisms of inflammatory resolution that al-low microglia to counterbalance activation signals. We identify the upregulation of exosome release as an important effector mechanism that promotes immune homeostasis in acutely activated and aged microglia. We show that microglia upregulate Rab27a-dependent exosome release during resolution of interferon-induced activation, a process that is partly mediated by anti-inflammatory cytokine signaling. These findings are in line with results showing increased exosome release in interleukin-4 treated bone marrow derived macrophages (Squadrito et al., 2014). Additionally, exosomes released by resolving microglia contain higher levels of immune proteins and miRNAs involved in toll-like and interferon signaling, such as miR-155. miR-155 has been implicated in driving pro-inflammatory transcriptional signatures in macrophages and in aberrant microglia inflammatory activation in disease conditions (Butovsky et al., 2015; Jablonski et al., 2016). Our experiments show that exosome release is critical for regulating cellular levels and activity of miR-155 in resolving microglia. Reduction of exosome release leads to miR-155 accumulation in resolving microglia, and as result sustained activation of STAT1 signaling and impaired inflammatory resolution. Previous studies showing that multivesicular bodies (MVB), the site of exosome formation, associate with (Gibbings et al., 2009) and regulate (Lee et al., 2009) miRNA-induced silencing complexes (miRSC) support dysregulation of miRNA activity as a mechanism driving this aberrant resolution phenotype. Inhibition of MVB formation results in decreased miRSC activity, while blockage of MVB turnover leads to over-accumulation of miRSCs (Gibbings et al., 2009; Lee et al., 2009). Our study shows that Rab27a knockdown, which causes cytoplasmic accumulation of MVBs (Ostrowski et al., 2010), phenocopies the effects of blockage of MVB turnover on miRSC activity. It is also plausible that increased miR-155 sorting into exosomes is caused by the decrease in pro-inflammatory genes targets during inflammatory resolution, based on a recent study showing that miRNAs against lowly expressed transcripts are directed towards exosomal secretion (Squadrito et al., 2014). Nonetheless, this scenario still supports our observations, as intracellular miR-155 accumulation would result in aberrant regulation of both lowly and highly expressed target transcripts during resolution. While our study focuses on miR-155, we do not exclude the role of other miRNAs and proteins given the heterogeneity in microglia exosome cargo and their associated biological functions. Indeed, gene ontology analyses of microglia exosome proteins suggests their involvement in protein trafficking and this is supported by a recent study showing that loss of Rab27a alters NADPH oxidase trafficking in activated macrophages and microglia (Ejlerskov et al., 2012). Further studies are necessary to better elucidate the diversity of exosome regulatory mechanisms and their relevance to different aspects of microglia inflammation.

Finally, we show dramatic alteration of exosome biogenesis in the aged brain, characterized by increased microglia exosome release and interstitial exosome abundance. Similar to our findings in acutely activated microglia, inhibition of exosome biogenesis in aged microglia exacerbates inflammatory activation. Transcriptional profiling recently revealed stronger anti-inflammatory gene profiles in aged compared to young microglia (Hickman et al., 2013). Our findings suggest exosome release as a possible effector of such anti-inflammatory gene networks in aged microglia. More work is, however, needed to understand interactions between exosomes and other microglia homeostatic mechanisms (Deczkowska et al., 2018; Matcovitch-Natan et al., 2016). Additionally, understanding the effects of dramatic endolysosomal reorganization in aged microglia on exosome cargo sorting and activity will better inform functional manipulation of exosomes in aged microglia. In summary, our findings identify exosome release as a novel anti-inflammatory effector mechanism utilized by acute and chronically activated microglia to restore immune homeostasis, and demonstrate a cell-state dependent dynamic interaction between exosomes, miRNAs and inflammation in microglia.

## Materials and Methods

### Animal Models

The following mouse lines were used: C57BL/6 (The Jackson Laboratory), C57BL/6 aged mice (National Institutes of Aging), C3H/HeSnJ wild-type and ashen mutant mice (The Jackson Laboratory). All studies were done in male mice. The numbers of mice used to result in statistically significant differences was calculated using standard power calculations with α = 0.05 and a power of 0.8. Power and size were calculated based on with the respective tests, variability of the assays and inter-individual differences within groups using http://www.stat.uiowa.edu/~rlenth/Power/index.html. Mice were housed under specific pathogen-free conditions under a 12 h light-dark cycle and all animal handling and use was in accordance with institutional guidelines approved by the University of California San Francisco IACUC.

### Stereotaxic injections

Animals were placed in a stereotaxic frame and anesthetized with 2% isoflurane (2L/ min oxygen flow rate) delivered through an anesthesia nose cone. Ophthalmic eye ointment (Puralube Vet Ointment, Dechra) was applied to the cornea to prevent desiccation during surgery. The area around the incision was trimmed. Solutions (GW4869 or DMSO dissolved in PBS) were injected unilaterally into the DG of the dorsal hippocampi using the following coordinates: (from bregma) anterior = -2mm, lateral = 1. 5mm, (from skull surface) height = -2.1mm. A 2.5 μl volume was injected stereotaxically over 10 minutes (injection speed: 0.20μl/min) using a 5 μl 26s gauge Hamilton syringe. To limit reflux along the injection track, the needle was maintained in situ for four minutes, slowly pulled out half way and kept in position for an additional two minutes. The skin was closed using silk suture. Each mouse was injected subcutaneously with the analgesic Buprenex. Mice were single-housed and monitored during recovery. Mice were perfused 7 days post-injection for tissue analyses.

### Intracerebroventricular (ICV) delivery

Osmotic pumps were first attached to an L-shaped cannula via a 3.5 cm tubing according to manufacturer’s instructions (Model 1002, Brain infusion kit 3). The assembly was preincubated in normal saline at 37C for 48-72hrs to activate pump prior to surgical implantation. The assembly was surgically implanted into the left ventricle of each mouse using a stereotactic apparatus, using the following coordinates: (from bregma) anterior = -0.4mm, lateral = 1.0mm, (from skull surface) height = -2.5mm. The osmotic pumps were place under the skin of the left flank by making a subcutaneous pocket with a curved, blunt scissors. Cannula were secured by gluing to the skull surface, prior to stitching the skin back together. Pumps contained either 10ng/ml IFNγ dissolved in PBS or just PBS. Pumps were loaded with a maximum volume of 100μl and pumped at a rate of 6μl per day. Pumps were implanted for either 3 or 7 days, before mice were sacrificed and brains harvested for histology, RNA and protein analyses.

### Microglia culture and isolation

BV2 microglia culture. The BV2 microglia cell line was maintained in growth media – DM EM (Thermo Fisher) supplemented with 10% heat-inactivated fetal bovine serum (FBS, Hyclone) and 1% penicillin/streptomycin (Life Technologies) in HERAcell 150i incubators (Caisson Labs) at 37C with 5% CO2. BV2 microglia were serially passaged once plates reached 80-90% confluency.

### Postnatal primary microglia isolation and culture

Primary microglial cells were harvested from mouse pups at postnatal day 3–6 (P3–P6). Briefly, the brain cortices were isolated and minced. Tissues were dissociated in 0.25% Trypsin-EDTA for 20 min at 37°C and agitated every 5 min. Trypsin was neutralized with complete medium (DMEM (Thermo Fisher) supplemented with 10% heat-inactivated fetal bovine serum (FBS, Hyclone)), and were filtered through 70μm cell strainers (BD Falcon) and pelleted by centrifugation at 1500 rpm. Mixed glial cultures were maintained in growth medium at 37°C and 5% CO2 for 7–10 d in vitro. Once bright round cells began to appear in the mixed glial cultures, recombinant mouse granulocyte macrophage colony stimulating factor (1ng/ml, Life Technologies) was added to promote microglia proliferation. Primary microglial cells were harvested by mechanical agitation after 48-72 hours and plated on poly-L lysine coated t-75 flasks (Corning) in growth media and used for functional assays within 72 hours of purification. Functional assays were performed in serum free media.

### Adult primary mouse microglia isolation

Wild type mice (3 and 24 months, National Institutes of Aging) were transcardially perfused with PBS, brains gently minced, and dissociated into single cells using the Neural Tissue Dissociation Kit P (Miltenyl Biotec). Cells were washed with Hank’s Balanced Salt Solution and resuspended in PBS containing 0.5% FBS and 2mM EDTA (MACS buffer). Myelin removal was performed by incubation of the cell suspension with a ferric anti-myelin antibody and passed through a LS separation column placed in a magnetic holder (Miltenyl Biotec). Microglia enrichment was performed on the flow through cells by incubation with CD11b microbeads (Miltenyl Biotec), washed with MACS buffer and applied to a LS separation column placed in a magnetic holder (Miltenyl Biotec). The columns were removed from the magnetic field and labeled cells were flushed from the column with MACS buffer. Flow cytometry on the labeled cells confirmed ~95% microglia purity. Isolated microglia were immediately processed for RNA isolation without culture or frozen in RNA stabilization buffer and frozen (RNAlater-ICE, Ambion). Quantitative PCR was performed on the samples using Taqman primers for IL1β, TNFα, IL10 and Gapdh (Life Technologies).

### Exosome isolation and characterization

*Isolation from microglia culture media*. BV2 or primary microglia were grown to ~60-70% confluency, thoroughly washed with PBS, and incubated in macrophage serum free media (MSFM) with or without interferon gamma. MSFM was collected after overnight incubation and processed for exosome isolation by differential ultracentrifugation, as previously described(Thery et al., 2006). Briefly, media was spun for 300 x g for 10 min to pellet live cells. Dead cells and apoptotic bodies were removed by sequential spins of 2,000 x g and 10,000 x g, respectively. The supernatant was then passed through a 0.22μm syringe filter unit. Exosomes were pelleted finally pelleted by centrifuging at 100,000 x g for 90 mins. Pelleted exosomes were then washed with 30ml filtered PBS at 100,000 x g for 90 mins. Ultracentrifugation was done using SW 28 or Ti 70.1 rotors on a Beckmann Coulter Optima XPN-80 ultracentrifugation. For mass spectrometry and Taqman microRNA profiling, exosomes were further concentrated using total exosome isolation buffer (Life Technologies). *Isolation of interstitial brain exosome*: Mouse brains were processed for exosome isolation following published techniques(Perez-Gonzalez et al., 2012). Briefly, freshly perfused or frozen hemibrains were gently homogenized in dissociation buffer - Hibernate E (Brain-Bits Inc.) plus papain (Wormingthon) and incubated at 37C for 15 mins. Digestion was quenched by diluting in twice the volume of Hibernate E, filtered through a 40μm mesh to eliminate debris. Further filtration was performed using a 0.22μm syringe filter. Differential ultracentrifugation was performed as described above to isolate exosomes. To concentrate exosomes into a smaller volume, the resuspended pellet from differential ultracentrifugation was incubated with total exosome isolation buffer (Life Technologies) overnight. Samples were centrifuged at 10000 x g for 60 mins, resuspended in sterile PBS, and utilized for downstream applications or stored at -80C. Nanoparticle tracking. Size distribution of the purified exosomes was determined using nanoparticle tracking analysis (Malvern Inc.). To get consistent measurements on the nanoparticle tracking software, purified exosomes were diluted hundred-fold or more in cold, filtered PBS to obtain an ideal vesicle concentration range. 3-4 measurements of each sample were collected and concentration values were averaged. *Transmission electron microscopy*. Exosomes were fixed in EM grade formaldehyde immediately following isolation. Negative staining was performed on a small aliquot of the fixed exosomes, which were loaded on Formvar-carbon coated EM grids and negative staining was performed as previously described(Thery et al., 2006). Scanning electron microscopy allowed visualization of vesicle structure and size. *Western Blot*. Exosomes were lysed in RIPA buffer (500 mM Tris, pH 7.4, 150 mM NaCl, 0.5% sodium deoxycholate, 1% NP40, 0.1% SDS, and complete protease inhibitors; Roche) or exosome lysis buffer (Life Technologies). Lysates were stored at -80C or used immediately for protein analysis. Lysates were mixed with 4x NuPage LDS loading buffer (Invitrogen) and loaded on a 10% SDS polyacrylamide gradient gel (Invitrogen) and subsequently transferred onto a nitrocellulose membrane. The blots were blocked in 5% milk in Tris-Buffered Saline with Tween (TBST) and incubated with rat anti-CD63 (1:500; MBL; clone: R5G2), rat anti-CD9 (1:2000, Biosciences; clone: KMC8), and rat anti-LAMP2 (1:200, Santa Cruz; clone: M3/84). CD9 and CD63 were run in non-reduced conditions. Protein lysates from BV2 microglia were probed with rabbit anti-phospho Stat1 (Tyr701, Cell Signaling), mouse anti-tubulin (Proteintech), and mouse anti-Gapdh (Sigma). Horseradish peroxidase-conjugated secondary antibodies (1:5000, GE Healthcare; NA934) and an ECL kit (Biorad) were used to detect protein signals. Multiple exposures were taken to select images with appropriate exposure (Biorad Imager). Selected images were exported at 300dpi and quantified using ImageJ software (Version 1.46k). Exosome Western blots were normalized to total protein concentration. *Liquid chromatography mass Spectrometry*. Pooled exosomes were lysed and denatured in an aqueous solution of 50mM TEAB and 0.1% SDS (Sigma), Protein concentration was measured using a BCA assay, and 25-50mg of each sample placed in a speedvac at 60C. Samples were reduced in 500mM TECP for 1 hour at 60C and alkylated by addition of 10mM iodoacetamide for 15 mins at room temperature. Trypsin (spectrometry grade, Promega) was added to the solution at ratio of 1:25 and incubate overnight at 37C. Detergent removal was performed on the samples following manufacturer’s instructions (Thermo Fisher). Samples placed in a speedvac until nearly dry and re-suspended in to a concentration of 1 μg/μl in LCMS buffer A and stored at -80C. Peptides were separated using a nanoLC Ultra 2D Plus cHIPLC system (SCIEX) in serial two column mode with two nano cHIPLC columns (75 μm x 15 cm ChromXP C18-CL 3 μm 300 Å). The peptides were initially loaded onto the first column and washed with buffer A (2% acetonitrile/98% H2O/0.1% formic acid) for 30 min at a flow rate of 0.5 μL/min. The elution gradient was 2-30% buffer B (98% acetonitrile/2% H2O/0.1% formic acid) over 120 min at 300 nL/min. The TripleTOF 5600 equipped with a NanoSpray III source (SCIEX) was used for MS data acquisition. The IDA method was constructed to acquire a TOF MS survey scan at >30,000 resolution for 0.25 msec, followed by 20 MS/MS spectra in 3 s at >15,000 resolution with an exclusion time of 15 s. Protein identification was performed by using ProteinPilot v5.0 software (SCIEX) and the UniProt SwissProt v2011605 Mus musculus database using integrated false discovery rate analysis function with a concatenated reversed database. Data were searched in thorough mode with tryspin digestion and iodoacetamide modification. Proteins detected with local false discovery rate (FDR) ≤5% from each individual experiment were aligned to a search result from all samples searched together (Master) to create a comprehensive list of proteins and compare groups. The Master list for alignment included191 proteins detected with a global FDR ≤1%. From the Master list, only proteins detected in replicates within each group were considered for final comparisons. microRNA profiling. Exosomes were pooled from several experiments to have sufficient input material. Pooled samples of BV2 microglia exosomes lysed and processed for microRNA isolation using the miRNeasy kit according to manufacturer’s instructions (Qiagen). To measure miRNA composition, reverse transcription and preamplification was performed using Rodent miRNA primer pool A and B on 300ng of input RNA isolated from pooled activated and unactivated BV2 exosomes (Life Technologies). To measure the abundance of individual miRNA species, samples were loaded into Taqman Rodent Array microRNA A and B cards (version 3.0, Applied Biosystems) and quantitative PCR was performed. Resultant Ct values for each sample were normalized to U6 levels, and fold change was determined relative to untreated exosome miRNAs. miRNA expression in interferon gamma (IFNγ, Life Technologies) treated microglia was performed using taqman miRNA assays (Life Technologies) against mmu-miR-155, has-miR-125-5p, has-miR-103, snU6. Following reverse transcription, real time PCR was performed to measure expression of miRNAs across different samples. Expression was normalized to U6 levels.

### Immunohistochemistry

Tissue processing and immunohistochemistry was performed on free-floating sec-
tions following standard published techniques(Villeda et al., 2011). Briefly, mice were anesthetized with 400 mg/kg avertin (Sigma-Aldrich) and transcardially perfused with 0.9% saline. Brains were removed and fixed in phosphate-buffered 4% paraformaldehyde, pH 7.4, at 4C for 48 h before they were sunk through 30% sucrose for cryoprotection. Brains were then sectioned coronally at 40 μm with a cryomicrotome (Zeiss, Inc.) and stored in cryoprotective medium. Primary antibodies were: rabbit anti-Iba1 (1:2000, Dako), rat anti-CD63 (1:100, Santa Cruz) and rat anti-CD68 (1:200, Millipore). After overnight incubation, primary antibody staining was revealed using fluorescence conjugated secondary antibodies (Life Technologies).

### Viral Infection

BV2 microglia were maintained in DMEM media containing 10% FBS. Cells were transduced with lentivirus (LV) containing an shRNA plasmid targeting mouse rab27a or a scramble shRNA sequence using a previously published transduction protocol(Lucin et al., 2013). The lentivirus also expressed a copped GFP, to allow visualization of infected cells. BV2 were plated at a density of 1000 cells per well in 96-well plates, and infected at a multiplicity of infection of 50 or 100 in the presence of polybrene (8 μg/μl). After overnight infection, media was removed and replaced with normal growth media. Cells were allowed to grow for an additional 72 hours, split into 24 well dishes and treated with puromycin to select for infected cells. Selection was performed for 48 hours and surviving cells were assessed for GFP expression using the Accuri C6 machine. Knockdown was confirmed qPCR assessment of rab27a transcript levels, due to lack of a good commercial antibody. LV plasmids were prepared by performing lipofectamine transfection of 293T cells with vectors expressing desired transgene, Pax2, and VSVG. Following overnight incubation, transfection media was replaced with viral production media, DMEM plus viral boost reagent (Alstem). Virus production media was collected the following day, centrifuged to pellet cells and debris, and filtered through a 0.45 mesh (Maine Productions). Virus containing media was concentrated by high speed ultra-centriguation, resuspended in PBS, aliquoted, and used immediately for experiments or stored at -80C. Data and statistical analyses. All experiments were randomized and blinded by an independent researcher prior to pharmacological treatment or assessment of genetic mouse models. Researchers remained blinded throughout histological and biochemical assessments. Groups were un-blinded at the end of each experiment upon statistical analysis. Data are expressed as mean ± SEM. The distribution of data in each set of experiments was tested for normality using D’Agostino-Pearson omnibus test or Shapiro-Wilk test. No significant differences in variance between groups were detected using an F test. Statistical analysis was performed with Prism 5.0 software (GraphPad Software). Means between two groups were compared with two-tailed Student’s t test. Comparisons of means from multiple groups with each other or against one control group were analyzed with 1-way ANOVA followed by appropriate post-hoc tests.

## Acknowledgments

This work was funded by NSF predoctoral fellowship (J.C.U), Society for Neuroscience Scholars Program (J.C.U), Glenn Foundation (S.A.V), NIA (R01 AG053382, R01 AG055797).

Supplemental material upload as separate pdf

